# *In vivo* cerebral microdialysis validation of the acute central glutamate response in a translational rat model of concussion combining force and rotation

**DOI:** 10.1101/432633

**Authors:** Ian Massé, Luc Moquin, Chloé Provost, Samuel Guay, Alain Gratton, Louis De Beaumont

**Affiliations:** Research Center, Hôpital du Sacré-Cœur de Montréal, 5400 Gouin Ouest Blvd, Montreal, Quebec, Canada, H4J 1C5; Research Center, Douglas Institute, 6875 LaSalle Blvd, Montreal, Quebec, Canada, H4H 1R3

**Keywords:** mild traumatic brain injury, concussion, head acceleration, cerebral microdialysis, rat

## Abstract

Concussions/mild traumatic brain injury (mTBI) represent a major public health concern due to persistent behavioral and neurological effects. The mechanisms by which concussions lead to such effects are partly attributable to an hyperacute indiscriminate glutamate release. Cerebral microdialysis studies in rodents reported a peak of extracellular glutamate 10 minutes after injury. Microdialysis has the advantage of being one of the few techniques allowing the quantification of neurotransmitters *in vivo* and at different time points following injury. In addition to the clear advantages afforded by microdialysis, the Wayne State weight-drop model induces an impact on the skull of a subject unrestrained by the fall of a weight. The latter model allows rapid acceleration and deceleration of the head and torso, an essential feature in human craniocerebral trauma and a factor that is missing from many existing animal concussion models. In the present study, we applied the Wayne State procedure and microdialysis to document, in awake rats, the acute changes in extracellular hippocampal glutamate and GABA levels resulting from concussive trauma. We studied the dorsal CA1 hippocampal region as it contains a high density of glutamatergic terminal and receptors, thus making it vulnerable to excitotoxic insult. Using HPLC, dialysate levels of hippocampal glutamate and GABA were measured in adult male Sprague-Dawley rats in 10 min increments for 60 min prior to, during and for 90 min following concussive trauma induced by the Wayne State weight-drop procedure. Sham control animals were treated in the same manner but without receiving the concussive trauma procedure. Our results show that concussive trauma is followed, within 10 min, by a robust, transient 3-fold increase in hippocampal glutamate levels; such changes were not seen in controls. In contrast, GABA levels were unaffected by the concussive trauma procedure. The findings derived from the approach used here are generally consistent with those of previous other studies. They also provide a crucial *in vivo* validation of the Wayne State procedure as a model with promising translational potential for pre-clinical studies on early therapeutic responses to concussion.

## Introduction

Traumatic brain injury (TBI) is a pathophysiological disruption of brain function induced by an external mechanical force. Concussions/mild traumatic brain injuries (mTBIs) are most prevalent, accounting for 70–90% of cases ^1^. Advances in translational neuroscience, combined with increased attention from the medical community, have provided insight into the mechanisms by which a concussion leads to post-traumatic neurological symptoms such as motor and cognitive impairments.

The acute functional disturbances after a concussion are mostly attributable to two distinct, yet interrelated pathophysiological mechanisms ^2,3^: (1) a primary brain injury and (2) a neurometabolic cascade often referred to as a secondary brain injury. The primary brain injury results from the rapid acceleration and deceleration of the head and torso which produces a compression of the brain tissues followed by a stretching of these same tissues during the backlash, shearing and stretching the axons ^4–6^. The neurometabolic cascade is the indirect cellular response to the concussion that occurs in the minutes and days following the primary brain injury.

Many researchers have documented the temporal coincidence of the resolution of short-lived acute concussion symptoms and the subsiding of the neurometabolic cascade ^7^. therefore making it a primary therapeutic target for concussions. Previous work suggests that glutamate, the main excitatory neurotransmitter of the central nervous system ^8^, plays two pivotal roles in the neurometabolic cascade of concussions ^7^. First, the immediate and indiscriminate release of glutamate following concussions sets off an excitotoxic response resulting in neuronal damage, cell death and dysfunction of the surviving neurons, in particular via overactivation of the glutamatergic receptor, N-methyl-D-aspartate (NMDA) receptors. Second, persistent disruptions of excitatory glutamatergic circuits are involved in motor and cognitive impairments persisting even decades post-concussion ^9^.

Excitotoxicity is the result of an imbalance between glutamate and γ-aminobutyric acid (GABA) ^9^, the main inhibitory neurotransmitter ^10^. Importantly, the balance between glutamate and GABA plays a key modulating role on neurological function. Pyramidal neurons, located in the cortex and hippocampus of mammals, as well as the mesencephalon, hypothalamic and cerebellar neurons, produce glutamate essential to excitatory signaling pathways ^11^. For its part, GABA is produced by interneurons modulating cortical and thalamocortical circuits, the latter circuits relaying sensory information and playing a key modulating role in the coordination of motor functions, attention and memory ^12^. GABA is also known to modulate the activity of excitatory pathways found throughout the brain and the loss of GABA-producing cells following concussions disrupts the equilibrium of excitation and inhibition leading to further cell damage and apoptosis ^9^. In addition, this excitatory imbalance in concussions was shown to accentuate cellular damage via diffuse axonal lesions and mitochondrial dysfunction ^7^.

The hippocampus, a brain structure heavily involved in cognitive function ^13, 14^, contains a high density of glutamatergic receptors, making it particularly vulnerable to excitotoxicity. Rodents models of concussion associated hippocampal damage to impairments in learning spatial memory and fear conditioning ^15, 16^.

Microdialysis is a minimally-invasive sampling technique allowing rapid, *in vivo* quantification of neurotransmitters such as glutamate and GABA at different time points following injury without sacrificing the animal. Studies in rodents, combining the microdialysis technique with different injury models, demonstrated the immediate rise in extracellular fluid (ECF) glutamate following mild or severe TBI ^17–20^. However, depending on TBI induction models and injury severity, variations exist as to the duration of that ECF glutamate peak extracted from the hippocampus, going from only the first 10 minutes ^19, 20^ up to a gradual return to baseline levels within a few hours ^17, 18^. In one microdialysis study using an open-skull weight-drop to model mild and severe TBI, the acute glutamate peak within 10 minutes returning to baseline levels within 20 minutes of injury induction was accompanied by a peak of ECF GABA, but the article did not display GABA data ^20^.

However, the inherent lack of ecological validity of concussion induction models such as the, controlled cortical impact and open-skull weight drop with respect to the biomechanics of concussion and the severity of induced brain damage hampers their translational value. Indeed, these procedures involve a craniotomy or a craniectomy, the head of the animal is therefore completely restricted in a stereotaxic apparatus, preventing the rapid acceleration and deceleration of the head and torso. Moreover, these models involve direct loading of the brain, inducing injuries significantly more severe than concussions suffered in humans.

In contrast, the mTBI induction model recently proposed by the Wayne State University ^21^ allows the induction of an impact to the skull (close as opposed to open-skull) of an animal not restricted by the fall of a weight, therefore allowing a rapid acceleration and deceleration of the head and torso. This acceleration/deceleration of the head and torso is a core biomechanical characteristic of concussions observed in humans that has failed to be addressed in previous animal concussion models. This method does not require incision of the scalp or surgery, and the procedure can be performed in less than a minute. Rodents were shown to recover spontaneously the righting reflex and show no signs of seizures, paralysis or altered behavior. Skull fractures and intracranial bleedings are very rare. Rodents show minor deficits in motor coordination. Histological analyzes reveal microglial activation and an increase in tau proteins. This animal model is simple, cost-effective and facilitates the characterization of the glutamate/GABA release arising immediately after a concussion.

### Objective

Given the alarming prevalence and the sometimes-catastrophic long-term consequences of concussions together with the intrinsic difficulties in developing a reliable and translational animal model of concussions, the aim of the current study was to develop a rat model of concussion, based on the Wayne State University model, which incorporates the microdialysis technique to study *in vivo* hyperacute extracellular glutamate and GABA changes over time following a concussion. Given the high density of glutamatergic receptors and its vulnerability to excitotoxic processes following concussions, extracellular glutamate and GABA will be measured from the hippocampus.

## Materials and Methods

### Animals

Male Sprague Dawley rats *(Charles River)* (n = 21) were delivered between 43 and 50 days of age and at a weight of 151–200 g (average 176 g). Rats were given 2 weeks to get used to their environment before the experimental protocol started. During these 2 weeks, the rats were handled for 5 minutes on a daily basis to facilitate their habituation in contact with the researchers. The rats were about 10 weeks old and weighing 295 to 351 g (mean 323 g) at the time of concussion induction. Throughout the protocol, the rats were housed in a cycle of 12 hours of light and 12 hours of darkness, at 24–26 degrees Celsius with continued access to water and food. A schematic outline of the research protocol is presented in figure 1. Cases details are summarized in table 1.

**TABLE 1:**
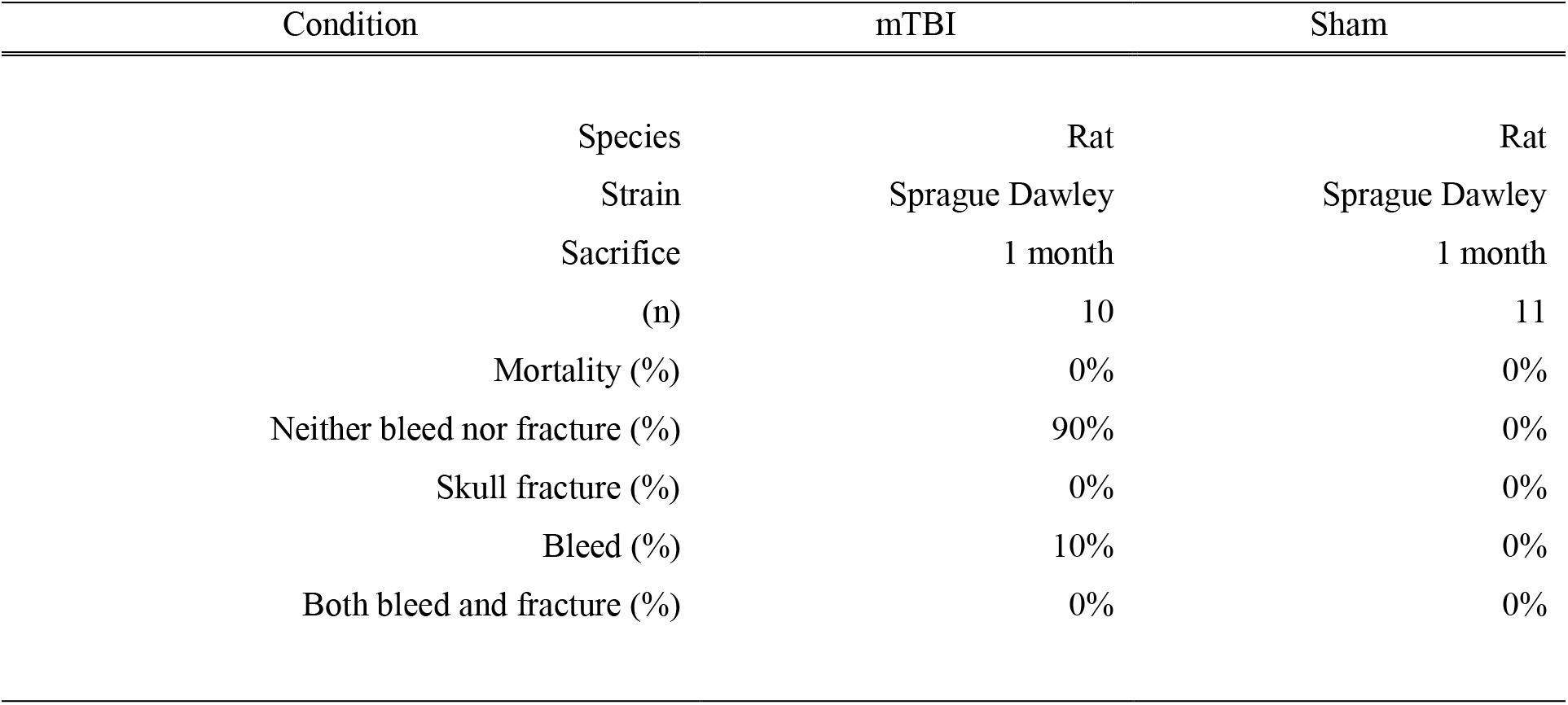
List of our cases analyzed by microdialysis (n = 21) and their general responsiveness.

**FIGURE 1:**
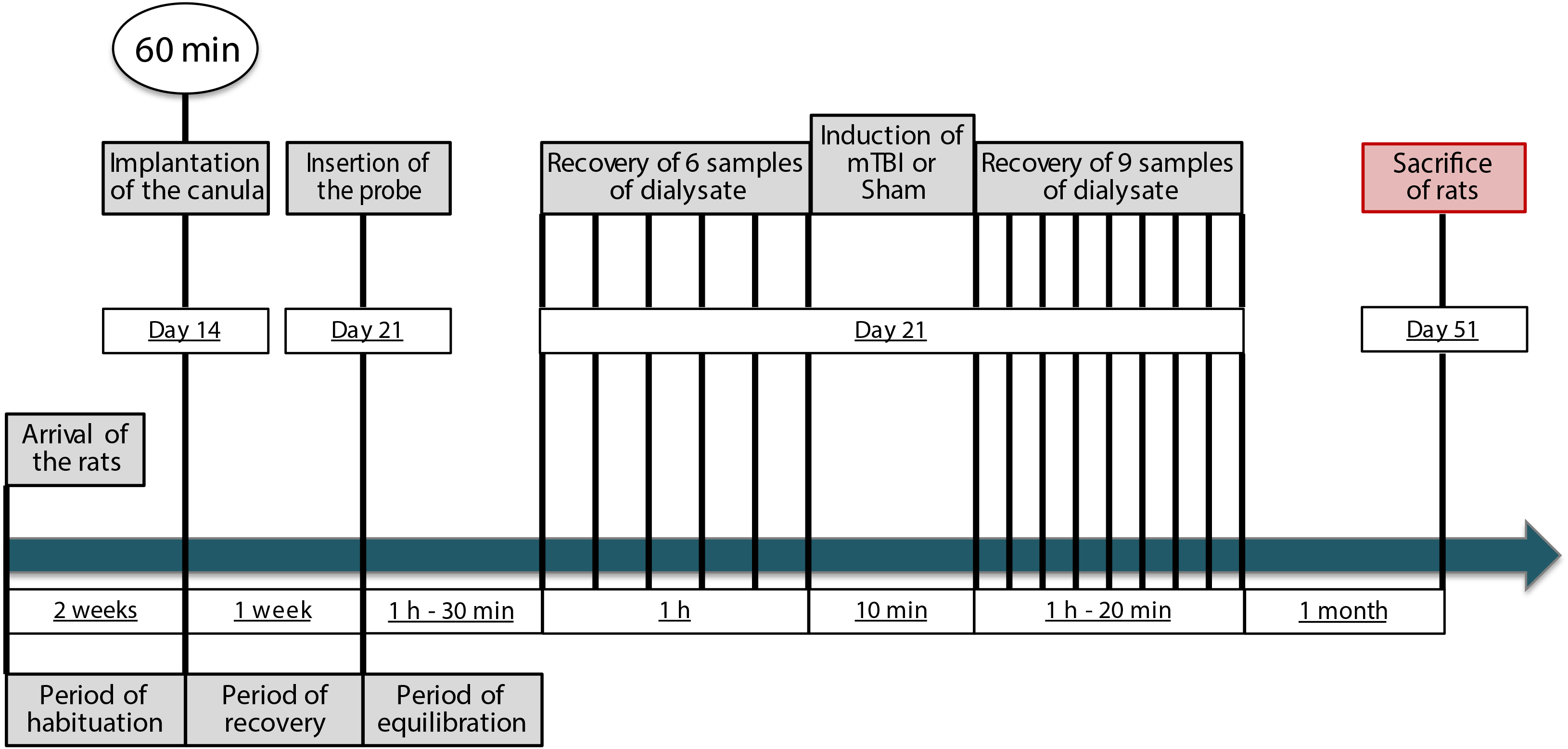
Schematic outline of the research protocol.

### Microdialysis guide cannula implantation surgery

Rats from the concussion group and the sham injury group were anesthetized with sodium isoflurane (2.5% isoflurane at 0.5 l/min oxygen flow), accompanied by infiltration analgesia of 1.5 mg/kg lidocaine/bupivacaine 10 minutes prior to incision of the skull and stereotaxically implanted with a 26-gauge stainless steel guide cannula *(Plastics One)* into the right hippocampus (AP: −6 mm, ML: −6.2 mm, DV: −1.6 mm, Fig. 2A) following Paxinos coordinates ^22^. These cannulas were used to insert the microdialysis probes into the target sites. Cannulas were secured with acrylic dental cement and 3 anchor screws threaded into the cranium. Buprenorphine (0.05 mg/kg, subcutaneously) was used for postoperative analgesia (once daily for 2 days). Animals were allowed 1 week (housed one per cage) to recover from cannula implantation before baseline and post-concussion measurements of extracellular glutamate and GABA levels by microdialysis. A removeable stainless steel obturator was inserted into the guide cannula to prevent CSF seepage and infection; the obturator was made to extent 2.5 mm beyond the tip of the guide cannula.

**FIGURE 2:**
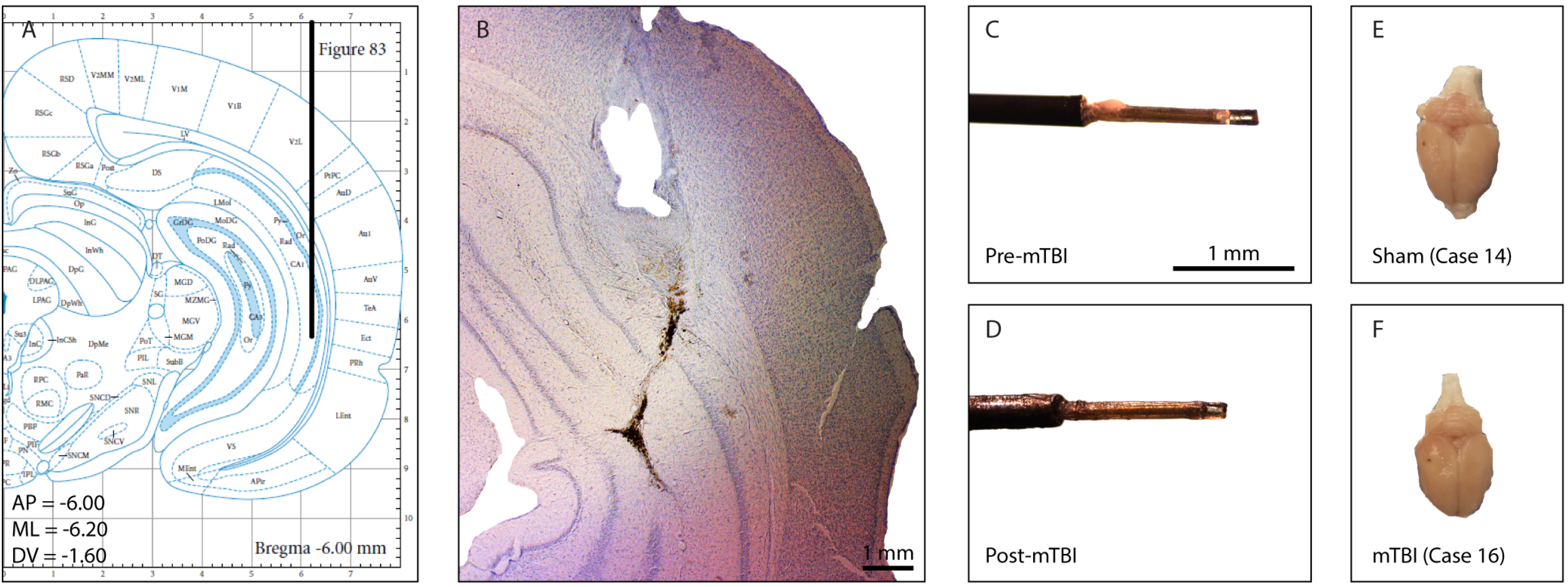
A: Coronal view of microdialysis probe and guide cannula placement site in the hippocampus using the stereotaxic atlas of Paxinos and Watson. B: Representative photomicrograph of hippocampus tissue damage (cresyl violet) produced by a microdialysis probe and guide cannula from a concussion case. C: Representative photomicrograph of a microdialysis probe before induction of concussion. D: Representative photomicrograph of a microdialysis probe after induction of concussion. The membrane is still intact. E-F: Representative photomicrograph of a sham (E) and concussion (F) injured brain following perfusion with 4% paraformaldehyde at 1-month after sham injury or concussion procedure. Upon visual inspection, the 2 brains are indistinguishable.

### Microdialysis probes

We used I-shaped, microdialysis probes comprised of side-by-side fused silica inlet-outlet lines [internal diameter (ID): 50 μm] that were encased in polyethylene tubing (ID: 0.58-0.38 mm). A length of regenerated, hollow cellulose membrane [Spectrum, molecular weight cut-off: 13 kDa, outer diameter (OD): 216 μm; ID: 200 μm] was secured to the end of a stainless-steel cannula (26-gauge) using cyanoacrylate adhesive and was sealed at its tip with epoxy; the active membrane measured 2.5 mm. A stainless-steel collar fitted to the probes provided a secure, threaded connection to the indwelling guide cannula of the animal. The probe assembly was affixed to a stainless-steel spring that was tethered to a liquid swivel *(CMA).* Probes were calibrated in artificial cerebrospinal fluid (ACSF) (26 mmol/l NaHCO3, 3mmol/l NaH2PO4, 1.3mmol/l MgCl2, 2.3mmol/l CaCl2, 3.0mmol/l KCl, 126mmol/l NaCl, 0.2mmol/l L-ascorbic acid). *In vitro* probe recovery ranged from 14% to 19% at a flow rate of 1 μl/min. A computer-controlled microinfusion pump (*CMA*) was used to deliver perfusate to the probes, and the dialysate was collected from the fused silica outlet line (dead volume: 0.79 μl).

### Microdialysis procedure

A microdialysis probe was inserted into the indwelling guide cannula of the unanesthetized animals and perfused with ACSF (flow rate set at 1 μl/min). Samples were then taken at 10-minute intervals for 60 minutes (baseline), followed by concussion (n = 10) or sham injury (n = 11), while still collecting samples for 90 minutes. Each 10- μl dialysate sample was collected in a fraction vial preloaded with 1 μl of 0.25 mol/l perchloric acid to prevent analyte degradation and immediately stored at 4°C for subsequent analysis.

### Concussion apparatus

The weights used to inflict the concussion (19 mm in diameter) were carved from solid brass to obtain the desired mass (450 g). The weights were dropped vertically through a PVC tube (20 mm diameter × 1.5 m length). The vertical trajectory of the falling weight was limited by a nylon fly fishing line (capacity of 9.1 kg, 0.46 mm diameter). A surface consisting of a slotted piece of aluminum foil held in place by a Plexiglas frame (38 cm long × 27 cm wide × 30 cm deep, Figs. 3A-B) held the rats in place. In this way, the slotted foil barely supported the body weight of a rat (295 to 351 g) with almost no resistance at impact. A foam cushion (37 cm long × 26 cm wide × 12 cm deep) was located beneath the aluminum foil leaf surface to cushion the falling rats while the weight remained attached over the free-falling body of the animals.

**FIGURE 3:**
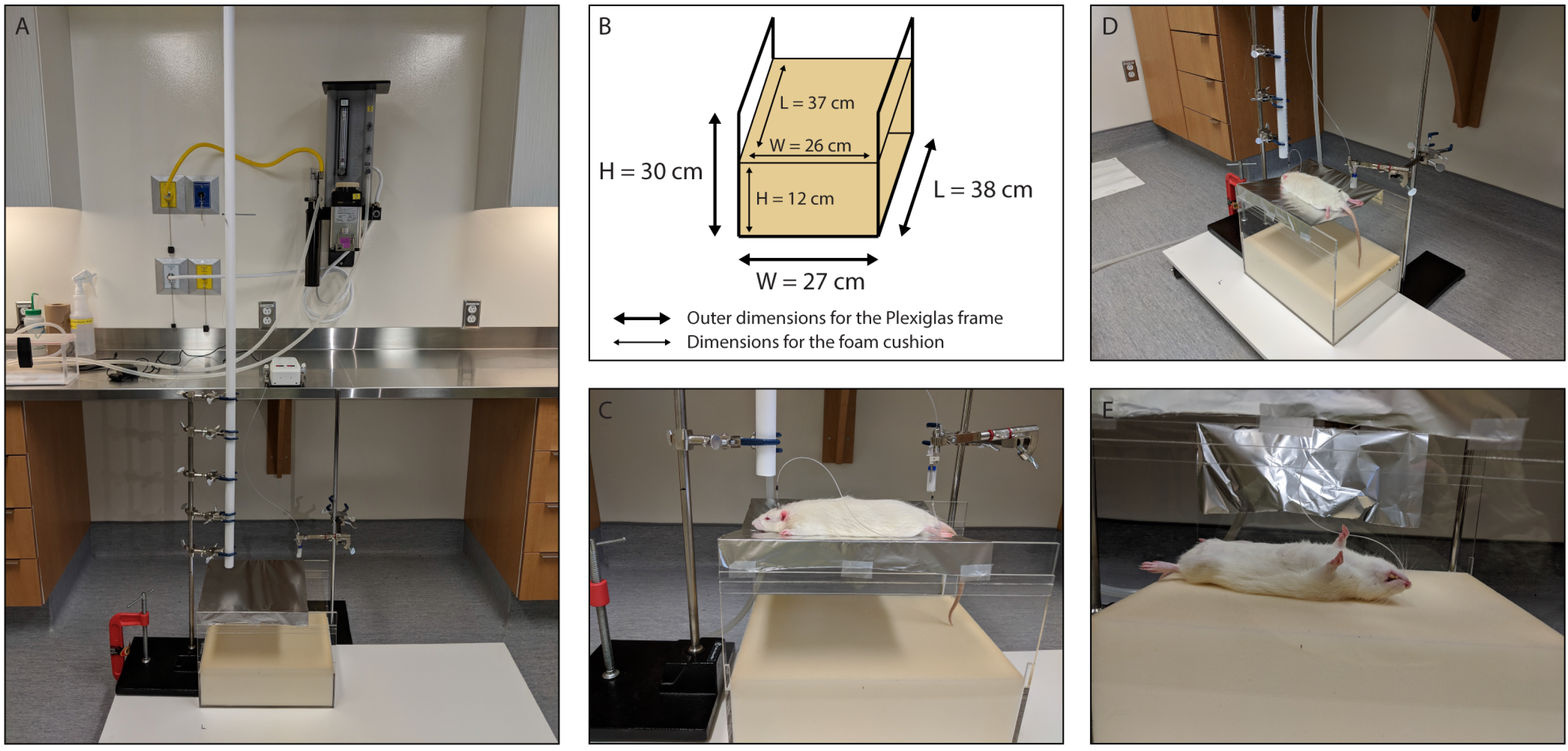
A-E: Concussion apparatus and microdialysis instruments essential components depictions. A: A photograph of the entire assembly comprised of a vertical polyvinyl chloride (PVC) guide tube for the falling weight situated above the rat stage, Plexiglas frame, foam cushion, computer-controlled microinfusion pump, gastight syringes, liquid swivels, and side-by-side fused silica inlet-outlet lines. B: Schematic representation of the Plexiglas frame and foam cushion with all pertinent dimensions. C: A photograph of the slotted piece of aluminum foil that serves as the rat stage above the foam cushion. D: A photograph showing the positioning of the rat on the stage immediately prior to head impact by the falling weight. E: A photograph showing the rat after head impact, illustrating the 180° horizontal rotation of the body of the rat after the head impact and ensuing acceleration and rotation.

### Induction of concussion and sham injury

The rats were slightly anesthetized with sodium isoflurane (until they no longer responded to the reflexes pinched to the paw or tail) and immediately placed under the PVC vertical tube. The rats were placed on the chest and supported by the piece of aluminum foil, over the foam cushion (Fig. 3C). The rats were rapidly positioned so that they were directly aligned with the weight by first placing the weight on the medial line of the scalp between the Bregma and Lambda sutures (Fig. 3D). The weight was then pulled up quickly by the line attached to the desired drop distance (1 m) and released. The descent of the falling weight was limited by the line so that after the first contact, the weight does not exceed by more than 2.5 cm the initial position of the dorsal surface of the head of the rats. Immediately after the impact, the rats fell on the foam cushion (Fig. 3E).

The acceleration and fall caused by the impact always involved a 180° horizontal rotation of the body of the rats. Subsequently, the rats were immediately moved into their cage to recover. Sham injury rats were also lightly anesthetized but without induction of trauma. A slow-motion video clip of the induction method for our combined rat model of concussion and cerebral microdialysis is included as Supplementary Material for online presentation.

### Righting time

Immediately following the induction of concussion or sham injury, rats were placed on their backs in their cage. Righting time was then acquired using a digital timer as an indicator of neurologic restoration. Righting time is the time taken by the rats to wake from the anesthetic and flip from the supine position to the prone position or begin walking. Any occurrence of bleed, fracture, or death was noted.

### High-performance liquid chromatography

Glutamate and GABA levels were determined as previously described by Lupinsky ^23^. A high-performance liquid chromatography precolumn derivatization with ultimate 3000 RS fluorescence detection (ex: 322 nm; emission: 455 nm) was used to determine glutamate and GABA levels. The chromatographic system consisted of a Dionex pump (ultimate 3000) and a Dionex RS autosampler (ultimate 3000) coupled to a Waters Xterra MS C18 3.0 × 50 mm 5 μm analytical column. The mobile phase was prepared as needed and consisted of 3.5% acetonitrile, 20% methanol, and 100 mmol/l sodium phosphate dibasic (Na2HPO4) adjusted to pH 6.7 with 85% phosphoric acid. The flow rate was set at 0.5 ml/min. Working standards (100 ng/ml) and derivatization reagents were prepared fresh daily from stock solutions and loaded with samples into a refrigerated (10°C) Dionex RS autosampler (ultimate 3000).

Before injection onto the analytical column, each fraction was sequentially mixed with 20 μl of o-phthaldehyde (0.0143 mol/l) diluted with 0.1 mol/l sodium tetraborate and 20 μl of 3-mercaptopropionic acid (0.071 mol/l) diluted with H2O and allowed to react for 10 minutes. After each injection, the injection loop was flushed with 20% methanol to prevent contamination of subsequent samples. Under these conditions, the retention time for glutamate and GABA was approximately 1 minute and 9.7 minutes, respectively, with a total run time of 30 minutes/sample. Chromatographic peak analysis was accomplished by identification of unknown peaks in a sample matched according to retention times from known standards using Chromaleon software. Analyte levels are expressed as μg/ml.

### Histology

One month after microdialysis and induction of concussion or sham injury, rats were anesthetized with ketamine/xylazine 100/10 mg/kg and then euthanized by intracardiac perfusion with saline and 4% paraformaldehyde. The rats were then decapitated and the brains dissected, stored in 4% paraformaldehyde and subsequently cryoprotected in a 30% sucrose solution. Brains were sliced in 50-μm-thick samples (coronal) and stained with cresyl violet (Nissl staining) for histological verification of probe placement and injury.

### Statistical analysis

Primary and secondary outcome continuous variables were analysed using Student t-tests. The Mann-Whitney U-test was used when data was not equally distributed. Normal distribution of continuous data was assessed with a Shapiro-Wilk test. All p-values were 2-tailed, and the significance level was set at 0.05. All analyses were performed using SPSS v 25.0 software (SPSS, Chicago, IL, USA).

## Results

### Histological verification of probe placement and injury

A total of 21 rats with histologically confirmed microdialysis probe placements in the CA1 region of the hippocampus were included in this study (concussion group, n = 10; sham injury group, n = 11). Histological examination of cresyl violet stained sections revealed no morphological change such as contusions or massive intracerebral hemorrhages due to concussion. Minimal and comparable damage between concussion and sham injury rats was due to implantation of the guide cannulas and insertion of the microdialysis probes. Induction of the concussion while a microdialysis probe was inserted did not produce distinguishable hippocampal tissue damage under the microscope (Fig. 2B). The membrane of the microdialysis probes remained intact after induction of the concussion procedure (Figs. 2C-D). No distinguishable difference was observed between the sham injury and concussion brains following perfusion with 4% paraformaldehyde at 1-month after microdialysis (Figs. 2E-F).

### Righting time

The time taken by rats to wake from the anesthetic and flip from the supine position to the prone position or begin walking following concussion or sham injury conditions is presented in figure 4. Rats from the concussion group took on average significantly longer to right themselves compared to the sham injury group (Student’s T-Test, p=0.042801). Although cases that experienced a concussion exhibited an increase in the righting time and appear stunned upon walking, they rapidly resume normal activities and were visually indistinguishable from sham injury cases. A single concussion case showed signs of minor bleeding beneath the site of impact, but did not show any sign of intracranial bleeding or skull fracture.

**FIGURE 4:**
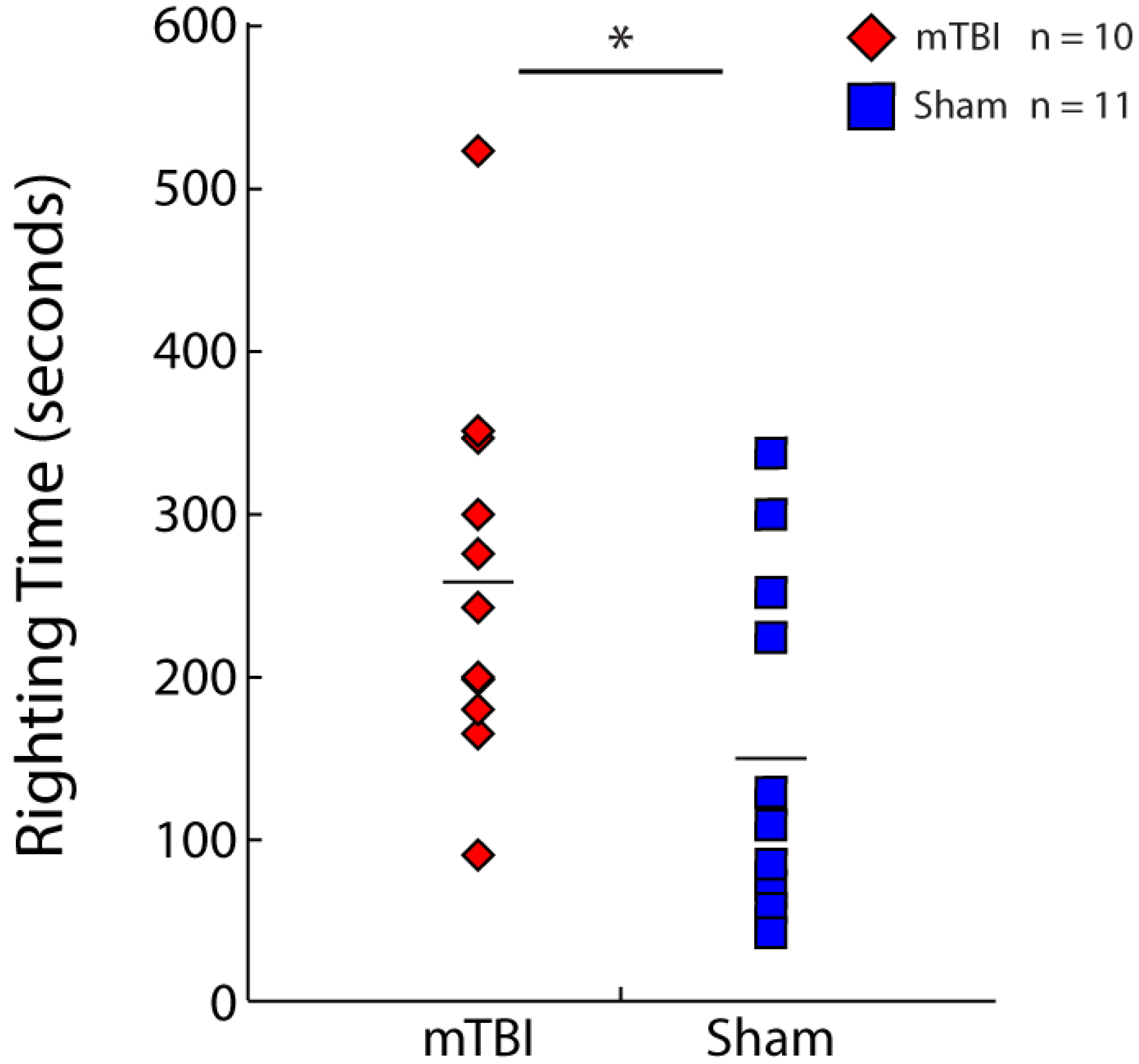
Histogram representations of the time taken by rats to wake from the anesthetic and flip from the supine position to the prone position or begin walking following concussion (red diamonds, n = 10) or sham injury (blue squares, n = 11) conditions. Rats from the concussion group took significantly longer to right themselves compared to the sham injury group. Mean values are represented as a horizontal line in each graph. * p<0.05, ** p<0.01, *** p<0.001.

### In vivo cerebral microdialysis

Extracellular concentrations of glutamate and GABA were analyzed from 15 in vivo, 10 μl dialysate samples obtained from the CA1 region of the hippocampus at 10-minute intervals with a flow rate of 1 μl/min during baseline (60 minutes/6 samples) and after concussion or sham injury (90 minutes/9 samples). Data on extracellular concentrations of glutamate and GABA for each individual case are available in table 2 and 3, respectively.

**TABLE 2:**
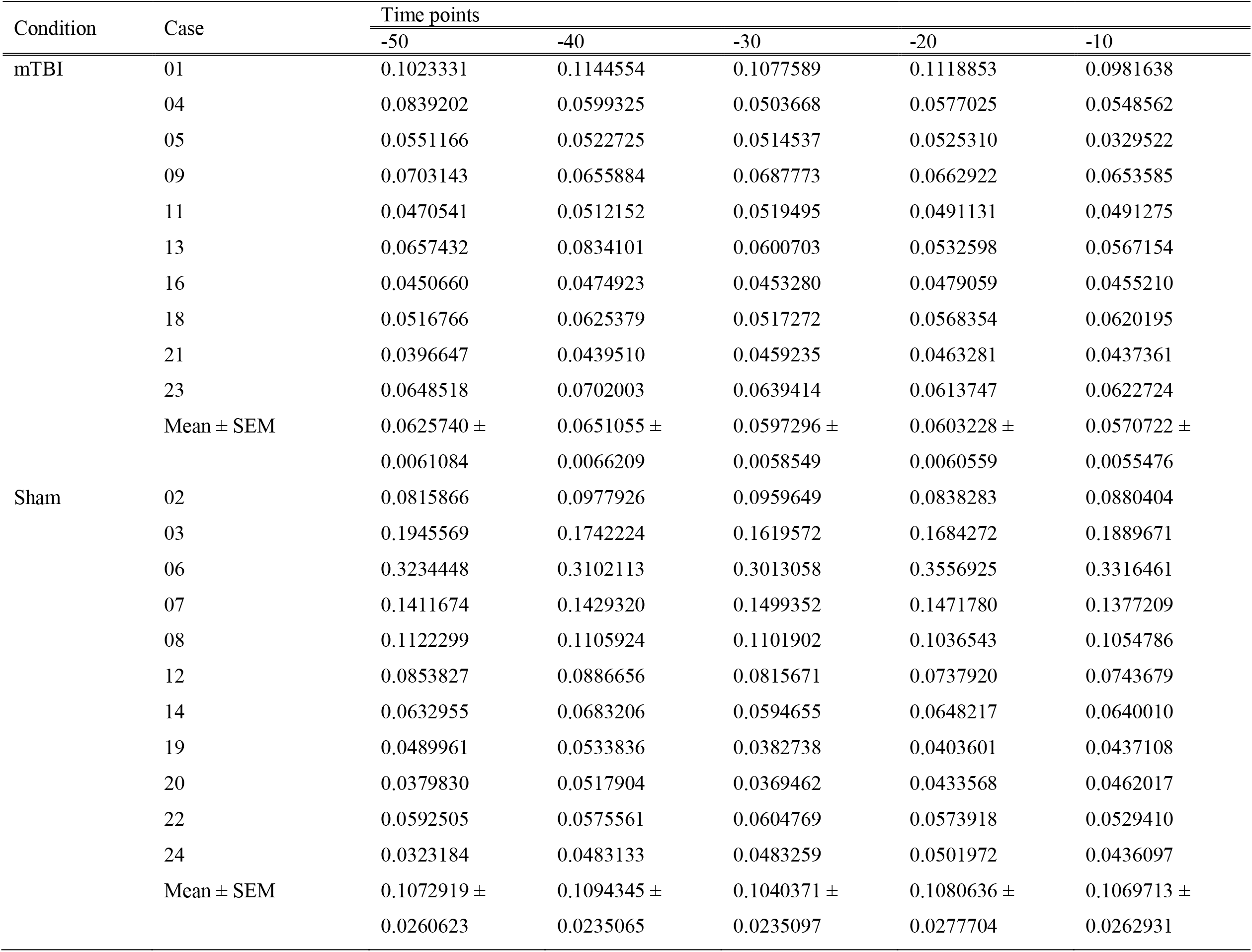
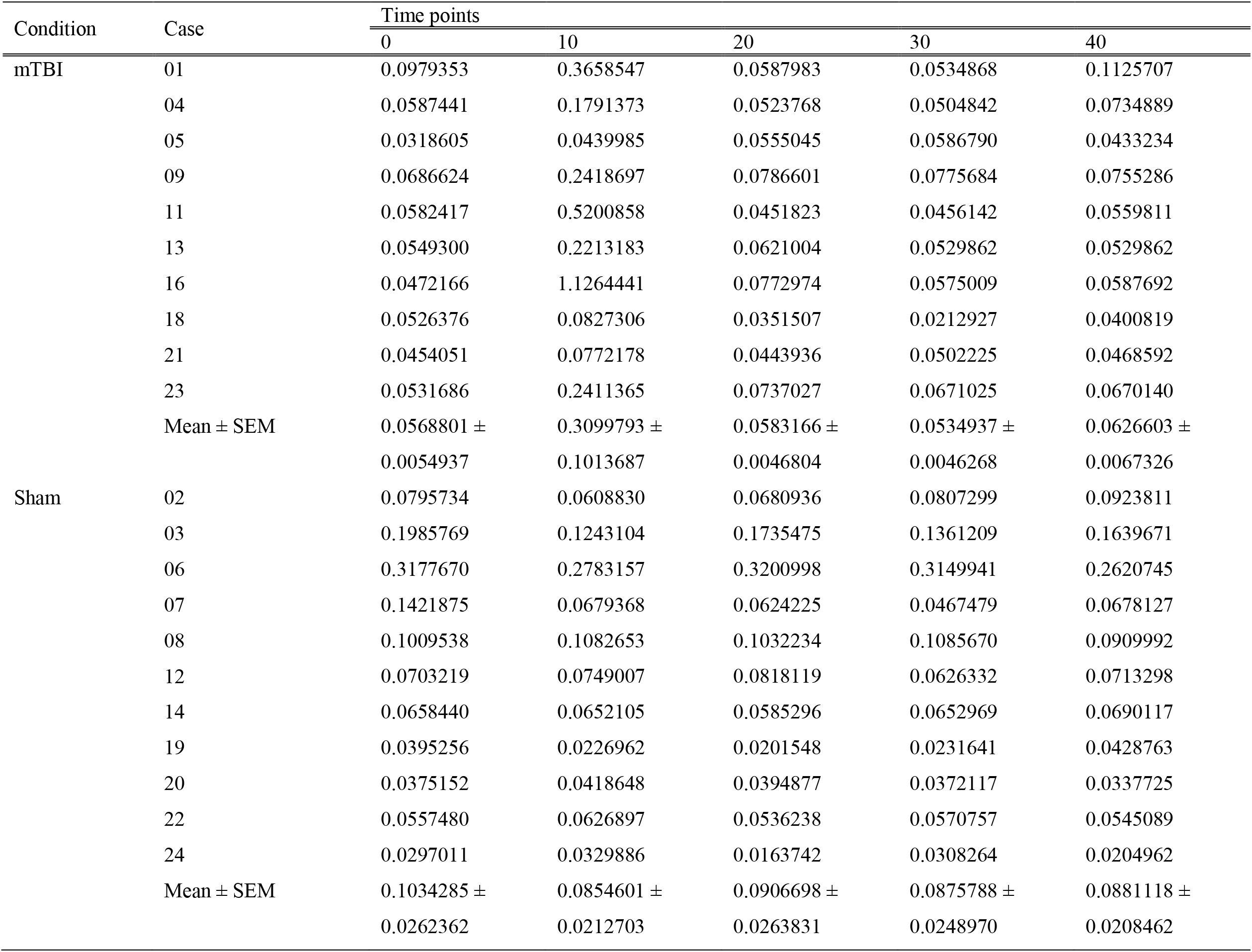
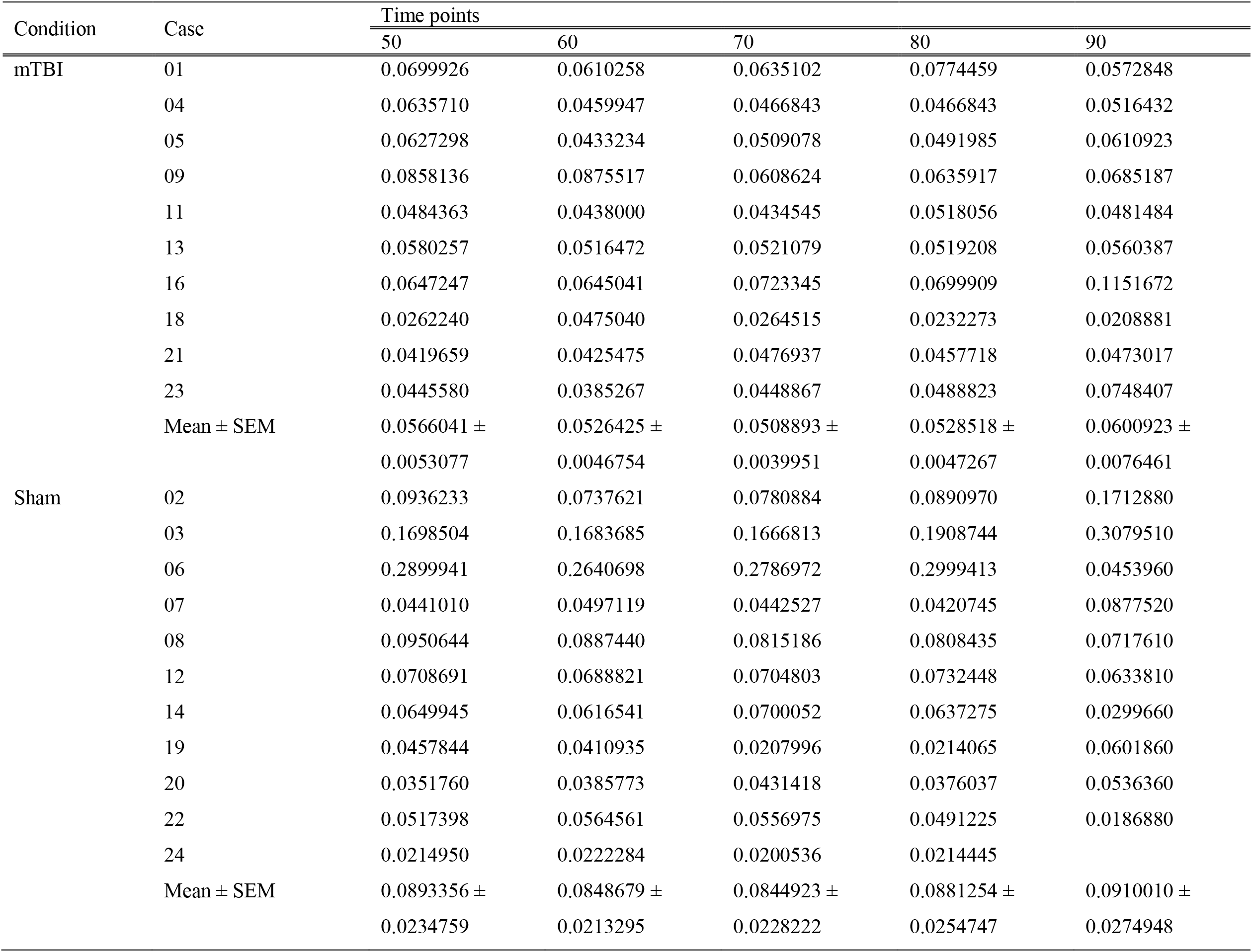
Extracellular concentrations of glutamate (ng/ml).

**TABLE 3:**
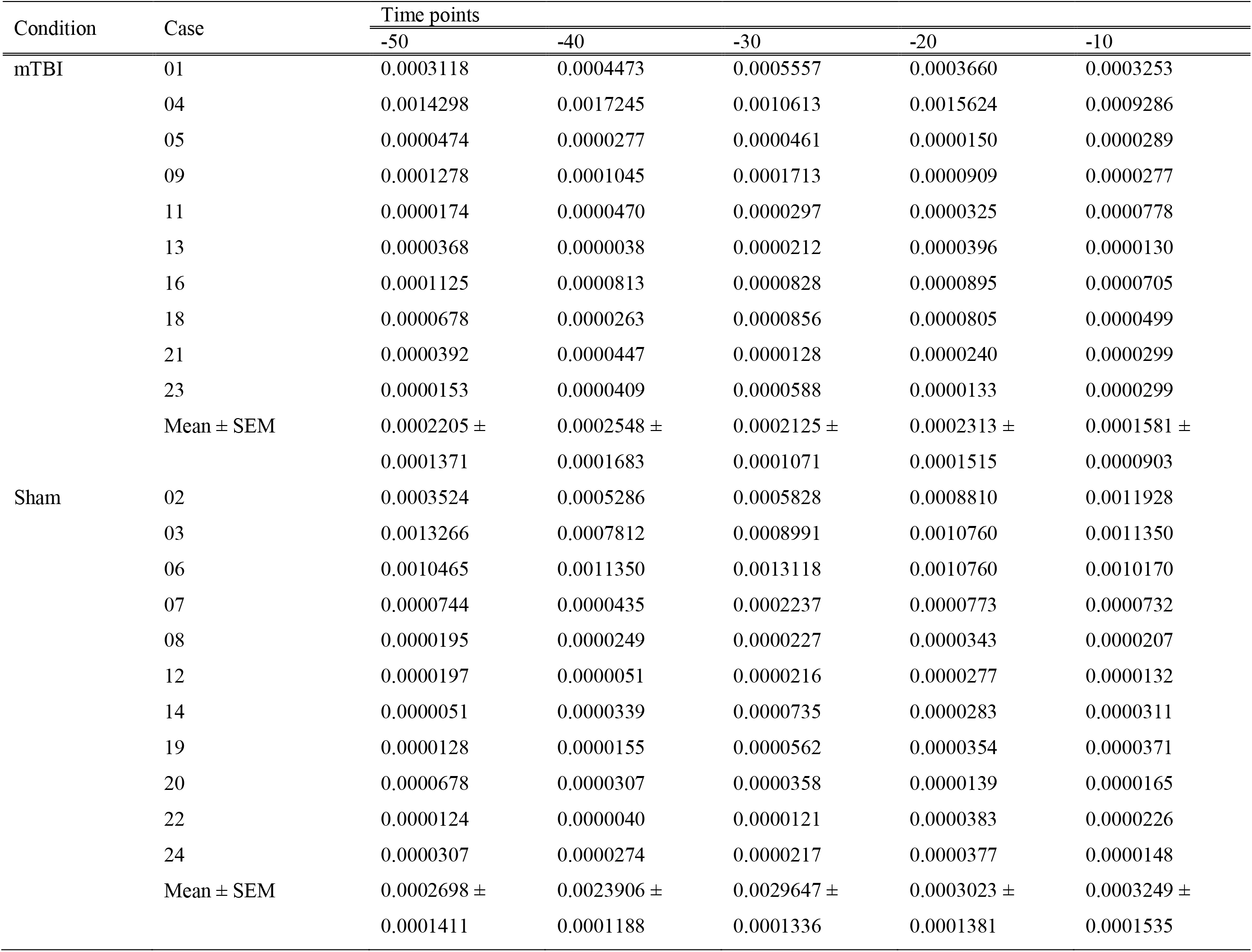
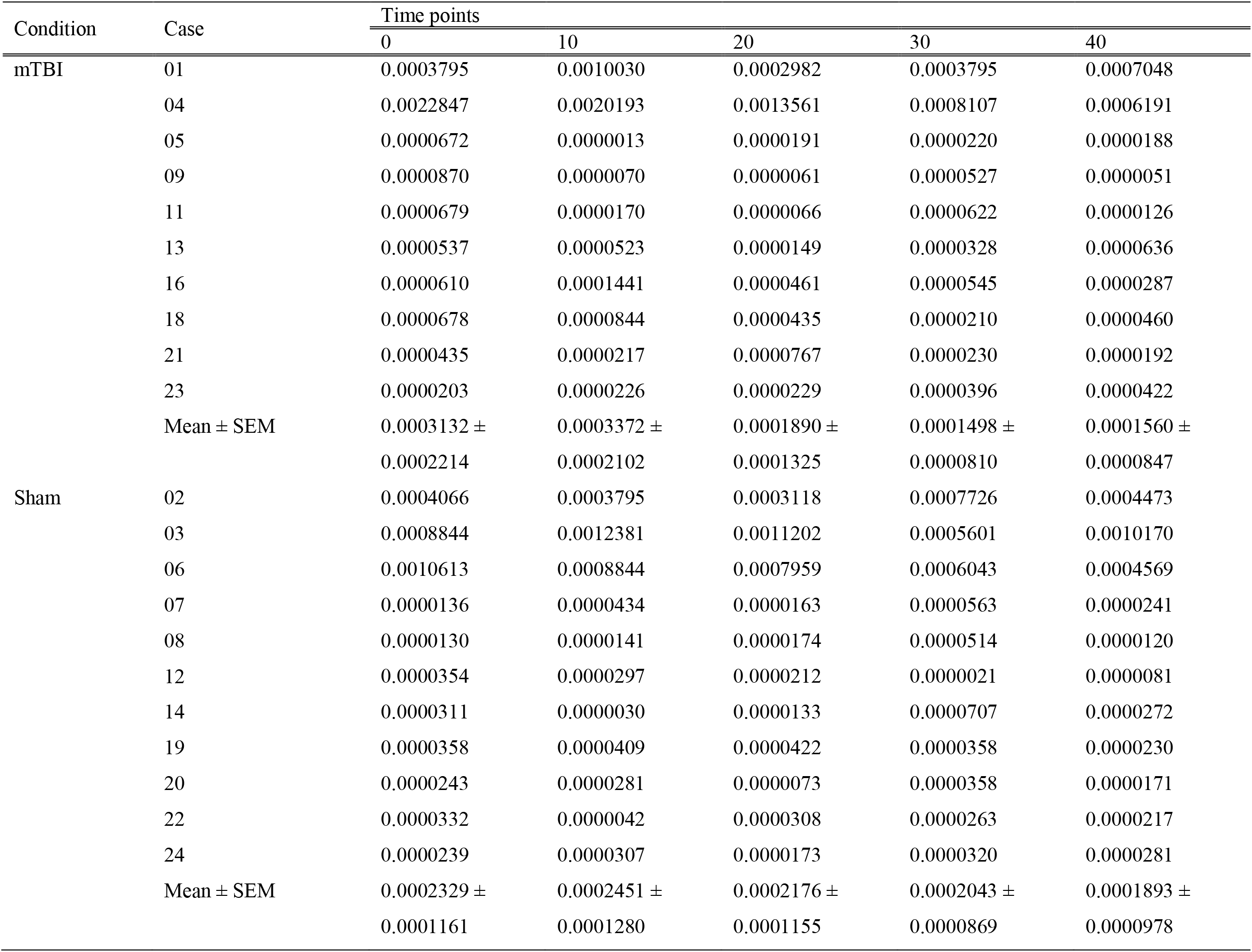
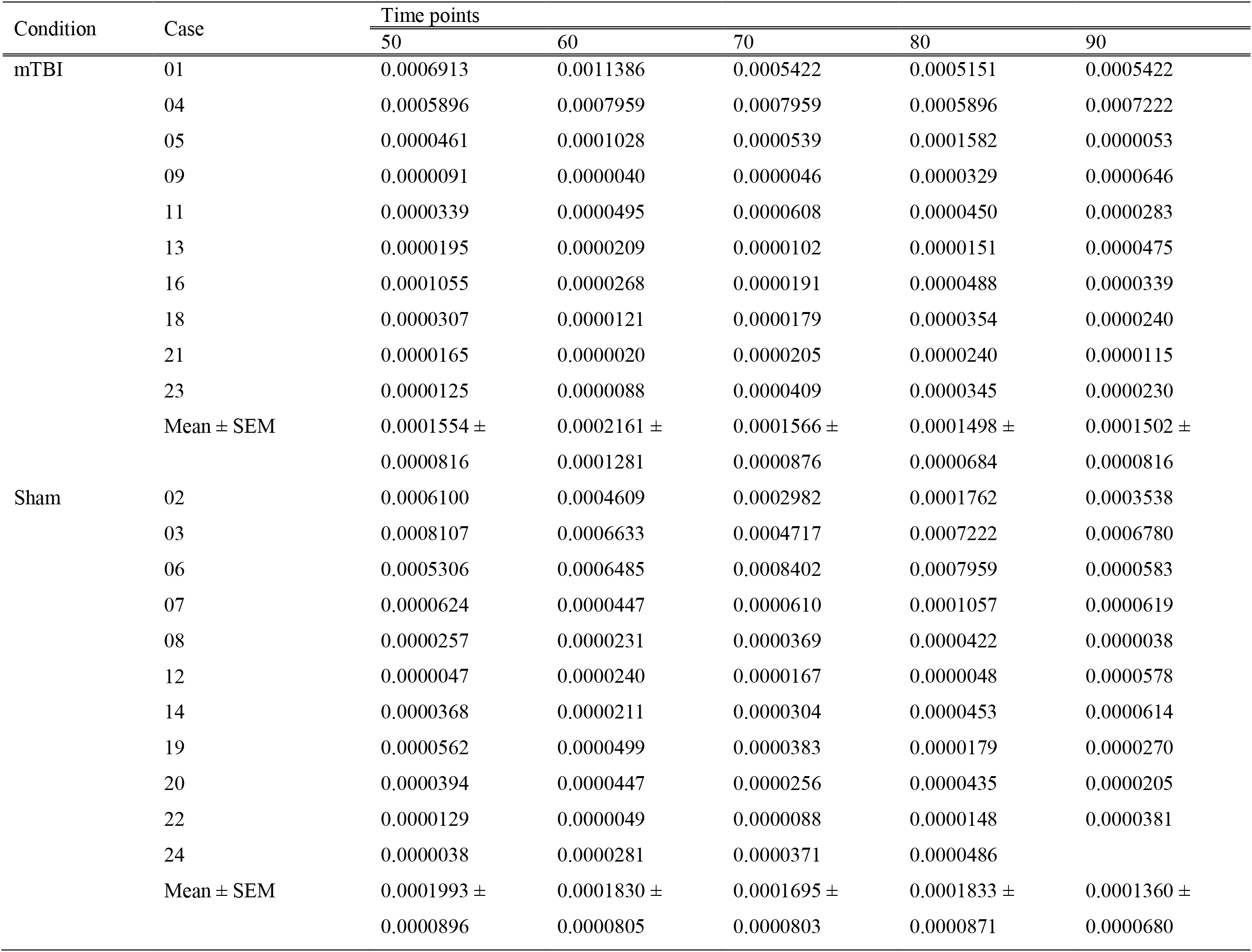
Extracellular concentrations of GABA (ng/ml).

### Extracellular concentrations of glutamate

Significant increases in glutamate levels were extracted from the hippocampus during the 10 minutes following concussion compared to sham injury (Mann-Whitney U Test, p=0.009175) (Fig. 5A). There was no other between-groups difference in glutamate levels at any other time points. The data are expressed in figure 5B and 5C for the concussion cases and sham injury cases, respectively, as the mean levels of glutamate during baseline and 10 minutes following concussion or sham injury. Baseline represents the averaged glutamate levels of the first 6 baseline samples collected before induction of concussion or sham injury. Relative to baseline levels, significant increases in glutamate levels were elicited in the hippocampus during the 10 minutes following concussion (Mann-Whitney U Test, p=0.004072) (Fig. 5B), but not following sham injury (Mann-Whitney U Test, p=0.450160) (Fig. 5C).

**FIGURE 5:**
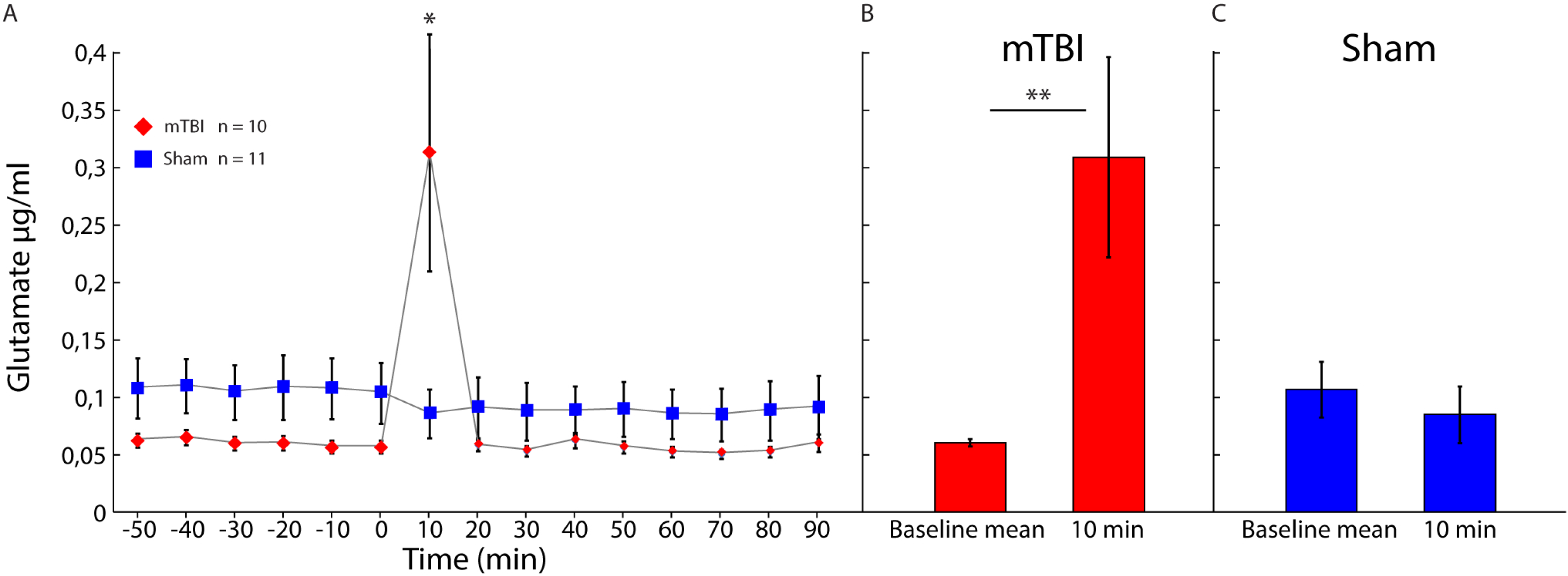
A: Mean extracellular concentrations of glutamate (ng/ml) measured by microdialysis in the hippocampus during baseline (60 minutes) and after concussion (red diamonds, n = 10) or sham injury (blue squares, n = 11) conditions (90 minutes). B-C: Comparison between mean levels of glutamate during baseline and 10 minutes following concussion (B) or sham injury (C). Error bars represent the standard error of mean. * P<0.05, ** P<0.01, *** P<0.001.

Data from extracellular concentrations of glutamate for each individual case in the concussion group and the sham injury group are represented in figure 6 and 7, respectively. Every case in the concussion group showed a peak increase in glutamate levels except for case 5 in which glutamate levels remained stable over the initial 10 minutes following concussion.

**FIGURE 6:**
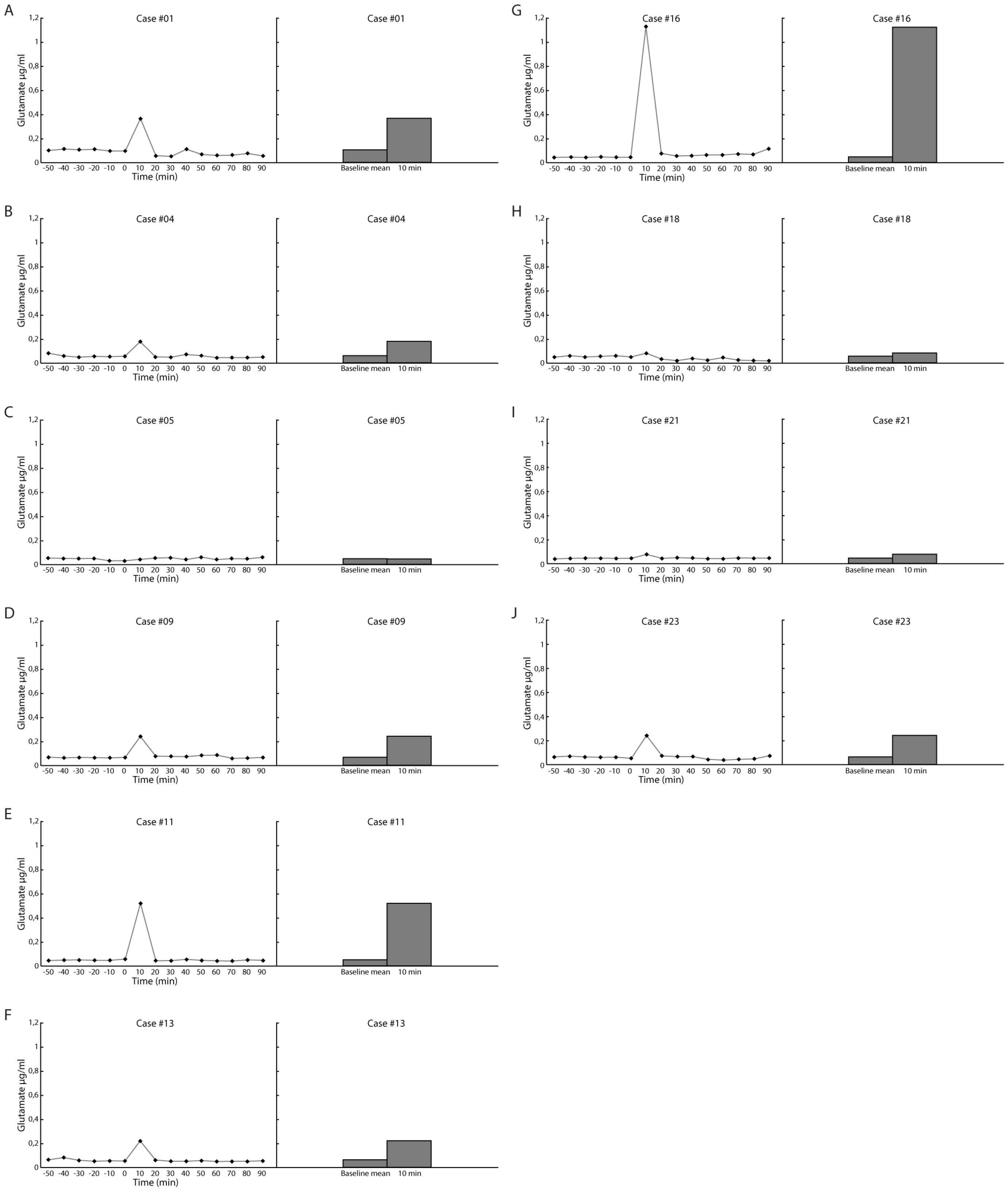
A-J: Extracellular concentrations of glutamate (ng/ml) measured by microdialysis in the hippocampus during baseline (60 minutes) and after concussion condition (90 minutes) for each individual case in the concussion group (curve). Comparison between mean levels of glutamate during baseline and 10 minutes following concussion for each individual case in the concussion group (histogram).

**FIGURE 7:**
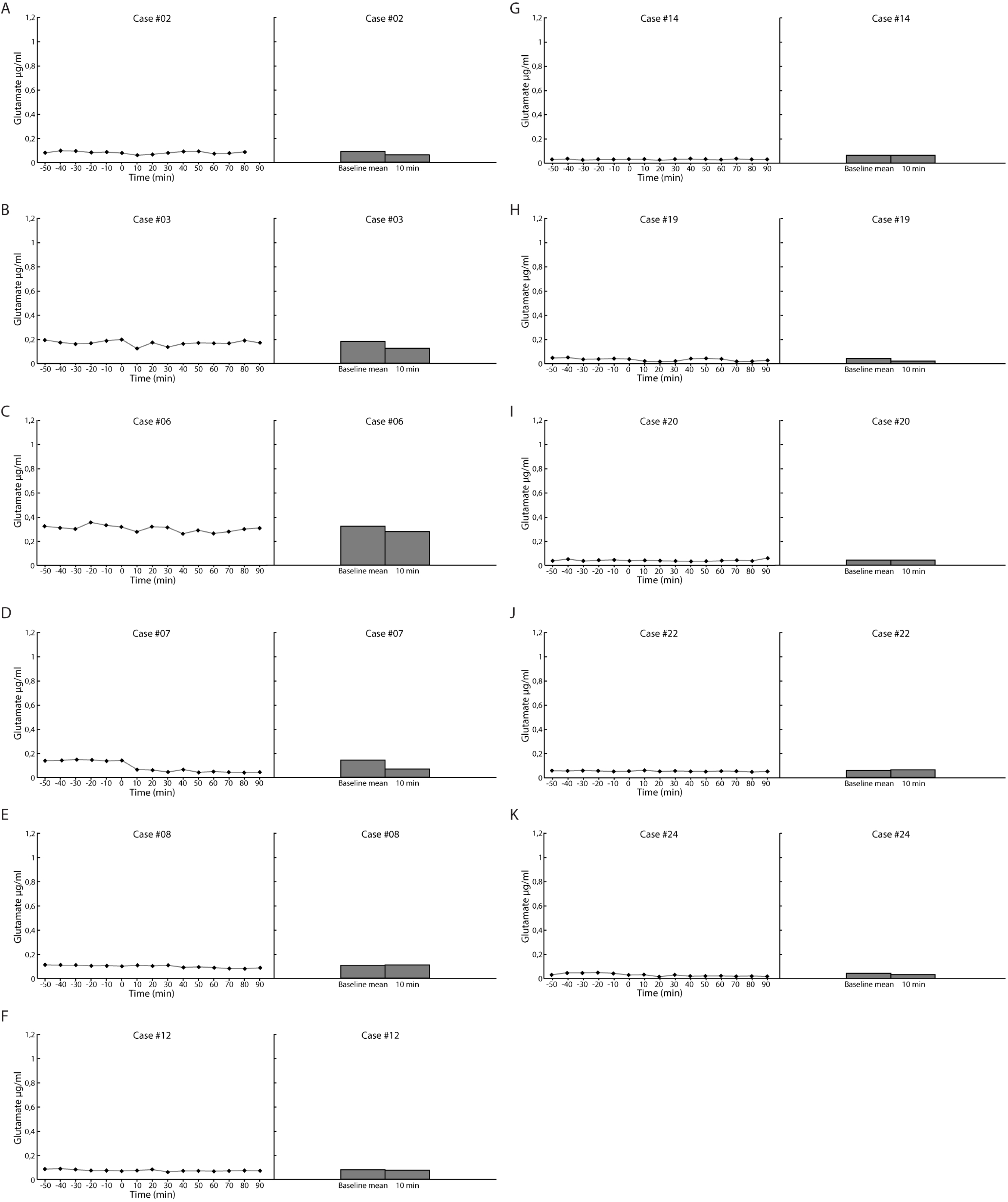
A-K: Extracellular concentrations of glutamate (ng/ml) measured by microdialysis in the hippocampus during baseline (60 minutes) and after sham injury condition (90 minutes) for each individual case in the sham injury group (curve). Comparison between mean levels of glutamate during baseline and 10 minutes following sham injury for each individual case in the sham injury group (histogram).

### Extracellular concentrations of GABA

There was no significant change in GABA levels extracted from the hippocampus during the 10 minutes following concussion compared to sham injury (Mann-Whitney U Test, p=0.943861) (Fig. 8). There was no other significant difference in GABA levels at any other time points between concussion cases and sham injury cases.

**FIGURE 8:**
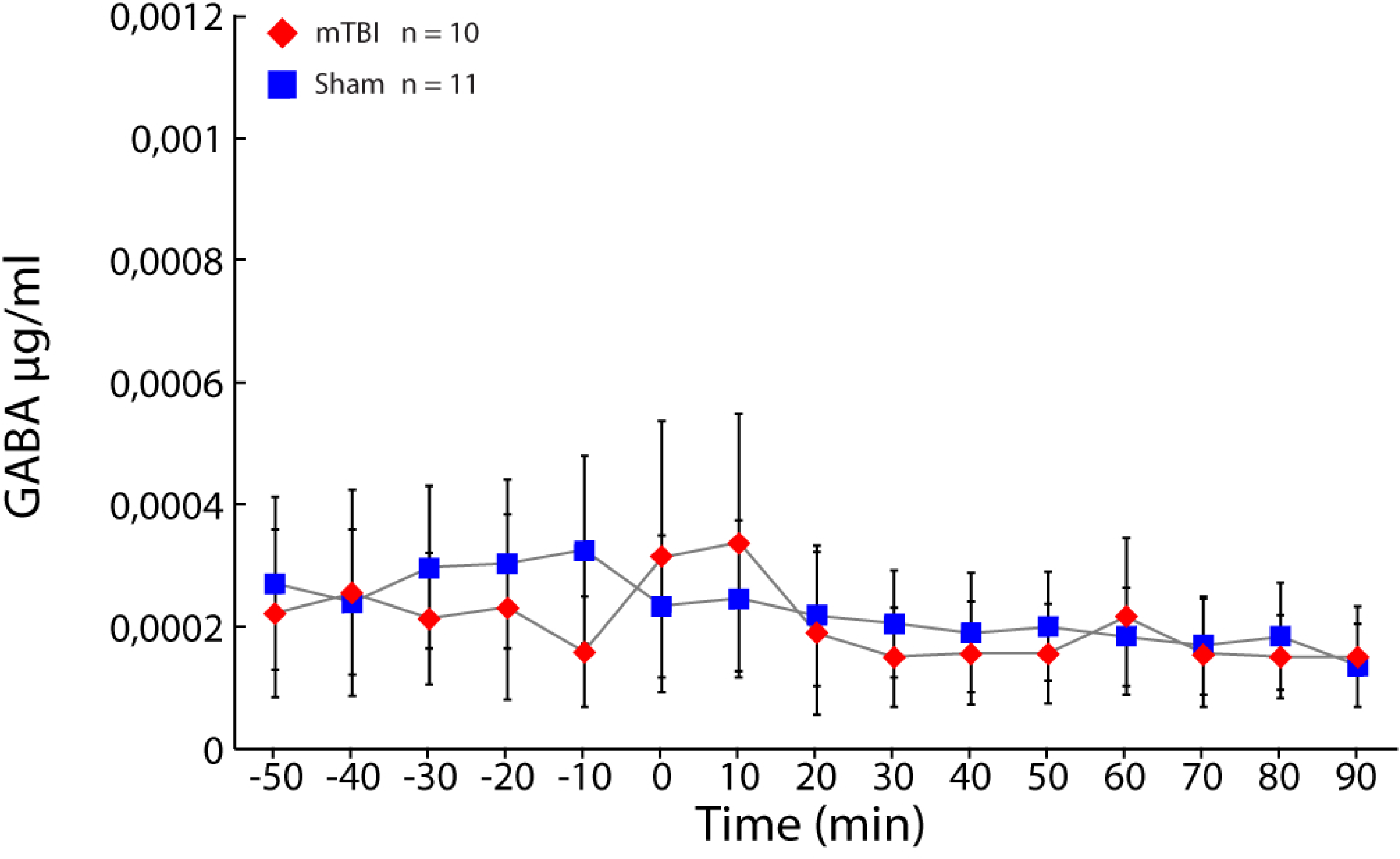
Mean extracellular concentrations of GABA (ng/ml) measured by microdialysis in the hippocampus during baseline (60 minutes) and after concussion (red diamonds, n = 10) or sham injury (blue squares, n = 11) conditions (90 minutes). Error bars represent the standard error of mean. * P<0.05, ** P<0.01, *** P<0.001.

Data from extracellular concentrations of GABA for each individual case in the concussion group and the sham injury group are represented in figure 9.

**FIGURE 9:**
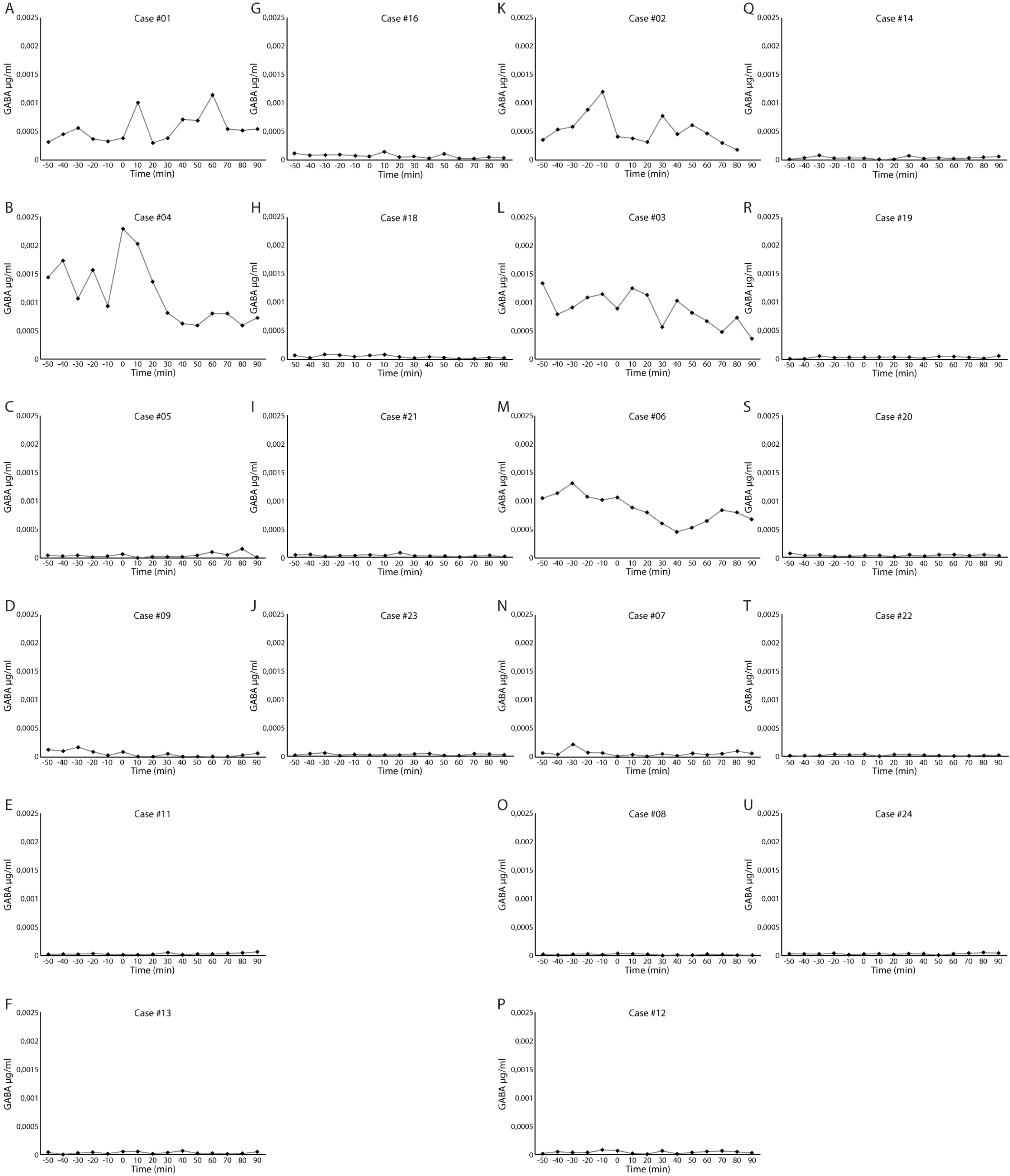
A-U: Extracellular concentrations of GABA (ng/ml) measured by microdialysis in the hippocampus during baseline (60 minutes) and after concussion (A-J) or sham injury (K-U) conditions (90 minutes) for each individual case.

## Discussion

### Main findings

This study aimed to develop a rat model of concussion, based on the Wayne State University model ^21^, which incorporates the microdialysis technique to investigate *in vivo* changes of extracellular glutamate and GABA over time. Given the high density of glutamatergic receptors and its vulnerability to excitotoxic processes following concussion, extracellular glutamate and GABA were measured from the hippocampus.

The main findings of this study are twofold: Firstly, we successfully induced a concussion using the Wayne State model while keeping the microdialysis probe inserted in the hippocampus throughout the entire experimental procedure. In all concussion cases (n=10), serious injury outcomes – namely high mortality, skull fracture, cardiorespiratory arrests and visible signs of cerebral contusions at the site of impact – were avoided. Secondly, our microdialysis procedure allowed us to replicate previous demonstrations of an hyperacute, short-lived extracellular glutamatergic release unfolding exclusively within the first 10 minutes of the impact ^17, 19^, However, in contrast with a previous microdialysis and TBI study in rats, our modified, close-skull weight-drop model did not reproduce the acute increase in extracellular GABA concentrations ^20^.

### Extracellular glutamate measured by microdialysis and methodological considerations

During the microdialysis experiment, peak ECF glutamate concentrations were found within the first 10 minutes following concussion induction, a finding that was exclusively observed in concussion rats (9 out of 10 concussion rats as opposed to 0 out of 11 sham rats). Over the subsequent 10 minutes – namely from 11 to 20 minutes following concussion induction – and onward for the following 70 minutes, glutamate concentrations were found to have returned to baseline levels. Our findings contrast with those of Faden et al. (1989) who had shown persistent glutamate concentration augmentations up to an hour following the injury. Well-documented differences in the severity of injury across animal models of TBI could at least partially account for study results discrepancies.

Given the known focal brain injury induced by probe insertion, the present microdialysis study included a sham group that underwent identical neurosurgical procedures but without subsequent concussion induction ^24^. Haemostatic and local environmental disturbances caused by probe insertion could result in temporary alterations of ECF glutamate concentrations ^24^. A previous study showed that even without inflicting any additional trauma, probes can give rise to a local immune response in the brain parenchyma up to two days following insertion ^24^.

Other than a slight, non-significant decrease observed within the first 10 minutes following induction of anesthesia with sodium isoflurane, ECF glutamate concentrations remained stable throughout the entire microdialysis experiment in sham cases. This suggests that the ECF glutamate concentration peak observed in rats from the concussion group was mostly due to the acceleration/deceleration of the head and torso following the weight-drop.

Leaving the microdialysis probe inserted in the hippocampus during impact is at odds with procedures described in most studies combining cerebral microdialysis and induction of TBI, which typically remove the probe during the insult and reposition it immediately after the injury ^19, 20, 25, 26^ in order to avoid damaging both the brain and the probe. However, the absence of any distinguishable difference in implanted hippocampal tissue visualized under cresyl violet-photomicrograph across concussion and sham groups as well as the stability of ECF GABA levels throughout all dialysate sampling times suggest that only slight, if any, additional damage to hippocampal tissue could be attributed to leaving the microdialysis probe inserted during impact ^27^. Alternatively, having kept the probe inserted throughout the entire experimental procedure significantly reduced the likelihood of inducing repeated damage associated with microdialysis probe insertion, including damage to the mTBI-sensitive blood-brain barrier ^28^. Interestingly, a one-time probe insertion into healthy brain tissue has been shown to leave the blood-brain barrier intact ^28^. Perhaps most importantly, uninterrupted dialysate collection throughout the entire experimental procedure allowed us to include a gold standard tissue recovery and equilibration period before beginning sample collection ^29^.

### Limitations of the current study and extracellular GABA measured by microdialysis

Although preliminary studies with the Wayne State model have assessed some basic molecular and structural changes ^21^, an in-depth analysis of the neuroanatomical and biological changes that occur at both cellular and epigenetic levels would reaffirm the validity of our model and its translational applicability. Moreover, assessment of cognitive dysfunction is a valid measure of outcome associated with concussion in animal models ^30^. Although we measured the righting time in this study and demonstrated significantly increased time in concussion cases relative to sham cases, future studies should focus on systematically evaluating cognitive performance after concussion induction in rats.

Furthermore, additional work is needed to clarify the potential role of extracellular GABA in concussion excitotoxicity. Our modified weight-drop model did not reproduce the increase in extracellular GABA concentrations reported in a previous microdialysis study ^20.^ Given the potential protective role of GABA against excitotoxicity following concussion, future studies should use microdialysis to systematically compare ECF GABA across TBI models and injury severities.

### Therapeutic applications

NMDA receptors antagonists are a very promising treatment perspective for conditions involving excitotoxicity such as concussion. In particular, the NMDA glutamatergic receptor antagonist, MK-801, has been shown to be an effective drug in decreasing hippocampus cell damage and preserving cognitive functions in rodents when injected immediately after concussion ^31, 32^. Although some protective effects of MK-801 have been demonstrated, its influence on the extracellular glutamate and GABA levels induced by the neurometabolic cascade following concussion is not yet fully understood. Important missing data are particularly related to the peak of extracellular glutamate reported within the first 10 minutes following the trauma ^17, 19^. The current concussion + microdialysis rodent model would be ideal in studying the acute ECF glutamatergic response to MK-801 or other NMDA receptors antagonist drugs.

## Conclusions

Our modified weight-drop model induced changes in ECF glutamate concentrations that are representative of the peak previously reported. Given the simple nature of the Wayne State model and the hyperacute ECF glutamate concentration changes measured using microdialysis, this combined rat model of concussion and microdialysis could provide researchers with a reliable and translational model of concussion that could allow longitudinal characterizations of the molecular effects of concussion.

## Acknowledgements

The authors report no conflict of interest. We are grateful to Louis Chiocchio for animal care and maintenance, Morgane Regniez for assistance with the intracardiac perfusion procedure, and David Castonguay for assistance with the cryostat. This work was supported by the Caroline Durand Foundation Chair in acute traumatology of the Université de Montréal awarded to LDB.

## Abbreviations

TBI: traumatic brain injury
mTBI: mild traumatic brain injury
ECF: extracellular fluid
CCI: controlled cortical impact
FPI: fluid percussion injury

## References

1. Cassidy, J.D., Carroll, L.J., Peloso, P.M., Borg, J., von Holst, H., Holm, L., Kraus, J., Coronado, V.G. and Injury, W.H.O.C.C.T.F.o.M.T.B. (2004). Incidence, risk factors and prevention of mild traumatic brain injury: results of the WHO Collaborating Centre Task Force on Mild Traumatic Brain Injury. J Rehabil Med, 28–60.

2. McCrory, P., Feddermann-Demont, N., Dvorak, J., Cassidy, J.D., McIntosh, A., Vos, P.E., Echemendia, R.J., Meeuwisse, W. and Tarnutzer, A.A. (2017). What is the definition of sports-related concussion: a systematic review. Br J Sports Med 51, 877–887.

3. McCrory, P., Meeuwisse, W.H., Dvorak, J., Echemendia, R.J., Engebretsen, L., Feddermann-Demont, N., McCrea, M., Makdissi, M., Patricios, J., Schneider, K.J. and Sills, A.K. (2017). 5th International Conference on Concussion in Sport (Berlin). Br J Sports Med 51, 837.

4. Cernak, I. (2005). Animal models of head trauma. NeuroRx 2, 410–422.

5. Davis, A.E. (2000). Mechanisms of traumatic brain injury: biomechanical, structural and cellular considerations. Crit Care Nurs Q 23, 1–13.

6. Gaetz, M. (2004). The neurophysiology of brain injury. Clin Neurophysiol 115, 4–18.

7. Giza, C.C. and Hovda, D.A. (2014). The new neurometabolic cascade of concussion. Neurosurgery 75 Suppl 4, S24–33.

8. Meldrum, B.S. (2000). Glutamate as a neurotransmitter in the brain: review of physiology and pathology. J Nutr 130, 1007S–1015S.

9. Guerriero, R.M., Giza, C.C. and Rotenberg, A. (2015). Glutamate and GABA imbalance following traumatic brain injury. Curr Neurol Neurosci Rep 15, 27.

10. Watanabe, M., Maemura, K., Kanbara, K., Tamayama, T. and Hayasaki, H. (2002). GABA and GABA receptors in the central nervous system and other organs. Int Rev Cytol 213, 1–47.

11. Spruston, N. (2008). Pyramidal neurons: dendritic structure and synaptic integration. Nat Rev Neurosci 9, 206–221.

12. Castro-Alamancos, M.A. and Connors, B.W. (1997). Thalamocortical synapses. Prog Neurobiol 51, 581–606.

13. Morris, R.G., Garrud, P., Rawlins, J.N. and O’Keefe, J. (1982). Place navigation impaired in rats with hippocampal lesions. Nature 297, 681–683.

14. Olton, D.S. and Papas, B.C. (1979). Spatial memory and hippocampal function. Neuropsychologia 17, 669–682.

15. Ray, S.K., Dixon, C.E. and Banik, N.L. (2002). Molecular mechanisms in the pathogenesis of traumatic brain injury. Histol Histopathol 17, 1137–1152.

16. Reger, M.L., Poulos, A.M., Buen, F., Giza, C.C., Hovda, D.A. and Fanselow, M.S. (2012). Concussive brain injury enhances fear learning and excitatory processes in the amygdala. Biol Psychiatry 71, 335–343.

17. Faden, A.I., Demediuk, P., Panter, S.S. and Vink, R. (1989). The role of excitatory amino acids and NMDA receptors in traumatic brain injury. Science 244, 798–800.

18. Folkersma, H., Foster Dingley, J.C., van Berckel, B.N., Rozemuller, A., Boellaard, R., Huisman, M.C., Lammertsma, A.A., Vandertop, W.P. and Molthoff, C.F. (2011). Increased cerebral (R)-[(11)C]PK11195 uptake and glutamate release in a rat model of traumatic brain injury: a longitudinal pilot study. J Neuroinflammation 8, 67.

19. Katayama, Y., Becker, D.P., Tamura, T. and Hovda, D.A. (1990). Massive increases in extracellular potassium and the indiscriminate release of glutamate following concussive brain injury. J Neurosurg 73, 889–900.

20. Nilsson, P., Hillered, L., Ponten, U. and Ungerstedt, U. (1990). Changes in cortical extracellular levels of energy-related metabolites and amino acids following concussive brain injury in rats. J Cereb Blood Flow Metab 10, 631–637.

21. Kane, M.J., Angoa-Perez, M., Briggs, D.I., Viano, D.C., Kreipke, C.W. and Kuhn, D.M. (2012). A mouse model of human repetitive mild traumatic brain injury. J Neurosci Methods 203, 41–49.

22. Paxinos, G. and Watson, C. (1998). The Rat Brain in Stereotaxic Coordinates. 4th ed. Academic Press: San Diego.

23. Lupinsky, D., Moquin, L. and Gratton, A. (2010). Interhemispheric regulation of the medial prefrontal cortical glutamate stress response in rats. J Neurosci 30, 7624–7633.

24. Woodroofe, M.N., Sarna, G.S., Wadhwa, M., Hayes, G.M., Loughlin, A.J., Tinker, A. and Cuzner, M.L. (1991). Detection of interleukin-1 and interleukin-6 in adult rat brain, following mechanical injury, by in vivo microdialysis: evidence of a role for microglia in cytokine production. J Neuroimmunol 33, 227–236.

25. Schwetye, K.E., Cirrito, J.R., Esparza, T.J., Mac Donald, C.L., Holtzman, D.M. and Brody, D.L. (2010). Traumatic brain injury reduces soluble extracellular amyloid-beta in mice: a methodologically novel combined microdialysis-controlled cortical impact study. Neurobiol Dis 40, 555–564.

26. Willie, J.T., Lim, M.M., Bennett, R.E., Azarion, A.A., Schwetye, K.E. and Brody, D.L. (2012). Controlled cortical impact traumatic brain injury acutely disrupts wakefulness and extracellular orexin dynamics as determined by intracerebral microdialysis in mice. J Neurotrauma 29, 1908–1921.

27. Chefer, V.I., Thompson, A.C., Zapata, A. and Shippenberg, T.S. (2009). Overview of brain microdialysis. Curr Protoc Neurosci Chapter 7, Unit7 1.

28. Sumbria, R.K., Klein, J. and Bickel, U. (2011). Acute depression of energy metabolism after microdialysis probe implantation is distinct from ischemia-induced changes in mouse brain. Neurochem Res 36, 109–116.

29. Zapata, A., Chefer, V.I. and Shippenberg, T.S. (2009). Microdialysis in rodents. Curr Protoc Neurosci Chapter 7, Unit7 2.

30. Bales, J.W., Wagner, A.K., Kline, A.E. and Dixon, C.E. (2009). Persistent cognitive dysfunction after traumatic brain injury: A dopamine hypothesis. Neurosci Biobehav Rev 33, 981–1003.

31. Han, R.Z., Hu, J.J., Weng, Y.C., Li, D.F. and Huang, Y. (2009). NMDA receptor antagonist MK-801 reduces neuronal damage and preserves learning and memory in a rat model of traumatic brain injury. Neurosci Bull 25, 367–375.

32. Sonmez, A., Sayin, O., Gurgen, S.G. and Calisir, M. (2015). Neuroprotective effects of MK-801 against traumatic brain injury in immature rats. Neurosci Lett 597, 137–142.

